# Selective Influence and Sequential Operations: A Research Strategy for Visual Search

**DOI:** 10.1101/619726

**Authors:** Kaleb A. Lowe, Thomas R. Reppert, Jeffrey D Schall

## Abstract

We introduce conceptually and empirically a powerful but underutilized experimental approach to dissect the cognitive processes supporting performance of a visual search task with factorial manipulations of singleton-distractor identifiability and stimulus-response cue discriminability. We show that systems factorial technology can distinguish processing architectures from the performance of macaque monkeys. This demonstration offers new opportunities to distinguish neural mechanisms through selective manipulation of visual encoding, search selection, rule encoding, and stimulus-response mapping.

## INTRODUCTION

To understand the neural mechanisms of visual search requires discovering the mapping between neural processes and visual, attention, and motor processes. Neural processes supporting visual search have been investigated in human studies using noninvasive measures of EEG and fMRI and in nonhuman primates using invasive sampling of neural discharges. Hence, to understand the neural mechanisms of visual search requires building a conceptual and empirical bridge between levels of explanation, neural measures, and species. Diverse computational and algorithmic approaches offer tools appropriate to translate between the neural and cognitive processes producing an observed pattern of performance. They serve another scientific function too. The literature on visual search and selective attention is governed by terms such as attention (both as cause and as effect), capacity, capture, disengage, efficiency, engage, map, priority, salience, selection, and shift. Formal models seem to us necessary to translate such common and vague psychological words into rigorous and useful scientific concepts.

### Visual search performance chronometry

Visual search takes time. A minimal amount of time is needed for visual encoding and response preparation. Not much more time is needed if the sought for object is easily discriminated from distracting objects, but progressively more time is needed if the distracting objects are more visually similar to the sought for target object and there are more such distracting objects (e.g, Treisman & Gelade 1980; Duncan & Humphreys 1989). Additional time may be taken is one of the non-target items is especially conspicuous (e.g., Theeuwes 1994; Bacon & Egeth 1994) or if the target item is in the same location as a previously attended target (Posner & Cohen 1984; Klein 2000). More time is needed if the response to the target object requires any kind of arbitrary mapping from stimulus location or property to response.

To investigate mechanisms of visual search at the neural circuit level requires systematic testing in nonhuman primates. For such studies to be relevant for understanding human performance, we must verify that nonhuman primates exhibit chronometric characteristics of search performance corresponding to humans. Fortunately, when sought, this confirmation has been found. Macaque monkeys exhibit dependence of visual search on target-distractor similarity and set size during singleton search (e.g., Azzato & Butter 1984; Buracas & Albright 1999; McPeek & Keller 2001; Sato et al. 2001; Arai et al. 2004; Camalier et al. 2007; Motter & Holsapple 2000, 2007; Balan et al. 2008; Song et al. 2008; Cohen et al. 2009; Nothdurft et al. 2009; Lee & McPeek 2013) and conjunction search (Motter & Belky 1998; Bichot & Schall 1999; Shen & Paré 2006). They can exhibit feature search asymmetries (Nakata et al. 2014). They can exhibit inhibition of return (Bichot & Schall, 2002; Fecteau & Munoz 2003; Shariat et al. 2012). Visual search is guided by memory as well as sensation. On the shortest time scale, performance of popout search varies if the search feature dimensions change (Malkovic & Nakayama, 1994). Called priming of popout, this demonstrated the limits of automaticity in visual search. Monkeys also exhibit priming of pop out (Bichot & Schall 2002). Macaque monkeys can also perform visual search filtering tasks that require search on one feature dimension and response according to another (Sato & Schall, 2003; Katnani & Gandhi 2013). Most recently, we have shown that monkeys also show contingent capture of attention by conspicuous non-target items (Cosman et al. 2018).

### Linking propositions through neural and mental chronometry

Understanding the neural mechanisms of visual search will need to explain the neural processes that occupy the different amounts of time. As visual search time increases, do a fixed number of neural processes just take longer? Or does an increase of visual search time happen because additional neural processes are inserted between encoding and responding?

A conceptually and historically foundational hypothesis posited that response time (RT) in complex tasks is the summation of functionally distinct stages (Donders 1868). The stage assumption is fundamental to the predominant model of “decision-making” – a single stochastic sequential-sampling process intervening between uninteresting visual encoding and response production stages (Ratcliff et al. 2016; Shadlen & Kiani 2013). Such models explain performance and account for neural activity in visual discrimination tasks as well as visual search with direct stimulus-response mapping (Purcell et al. 2010, 2012). But, if RT is not comprised of dissociable stages, then models like drift diffusion are disqualified and alternative models are endorsed, such as cascade (e.g., McClelland 1979) or asynchronous discrete flow (Miller 1988), which are qualitatively different mechanisms.

If mental modules are distinct and independent, then it should be possible to change one process without changing the other. The most effective method for assessing the existence of modules or stages is the logic of *separate modifiability* formulated by Saul Sternberg (2001). If mental modules are distinct and independent, then it should be possible to change one process without changing the other. The logical, mathematical, and statistical formulation developed by Sternberg specifies how to interpret the effects of specific causal manipulations on performance and neural measures. For example, if factors *F* (e.g, singleton-distractor identifiability) and *G* (stimulus-response cue discriminability) influence two sequential processes, A and B, selectively, then RT = Duration_**A**_(*F*) + Duration_**B**_(*G*). If A and B are distinct processes, then in an *F* × *G* factorial experiment, changes of RT over variation of *F* will be independent of changes of RT over variation of *G* (Figure 1). We can employ this logic systematically to determine which causal manipulations do and do not satisfy the criteria of additivity and mutual invariance. This approach has already revealed additivity and mutual invariance of singleton-distractor similarity and response interference in monkey cognitive neurophysiology studies (Moret & Hasbrouq 2000; Sato et al. 2001) and human ERP studies (e.g., Osman et al. 1992; Smulders et al. 1995; Servant et al. 2015).

**Figure 1.**
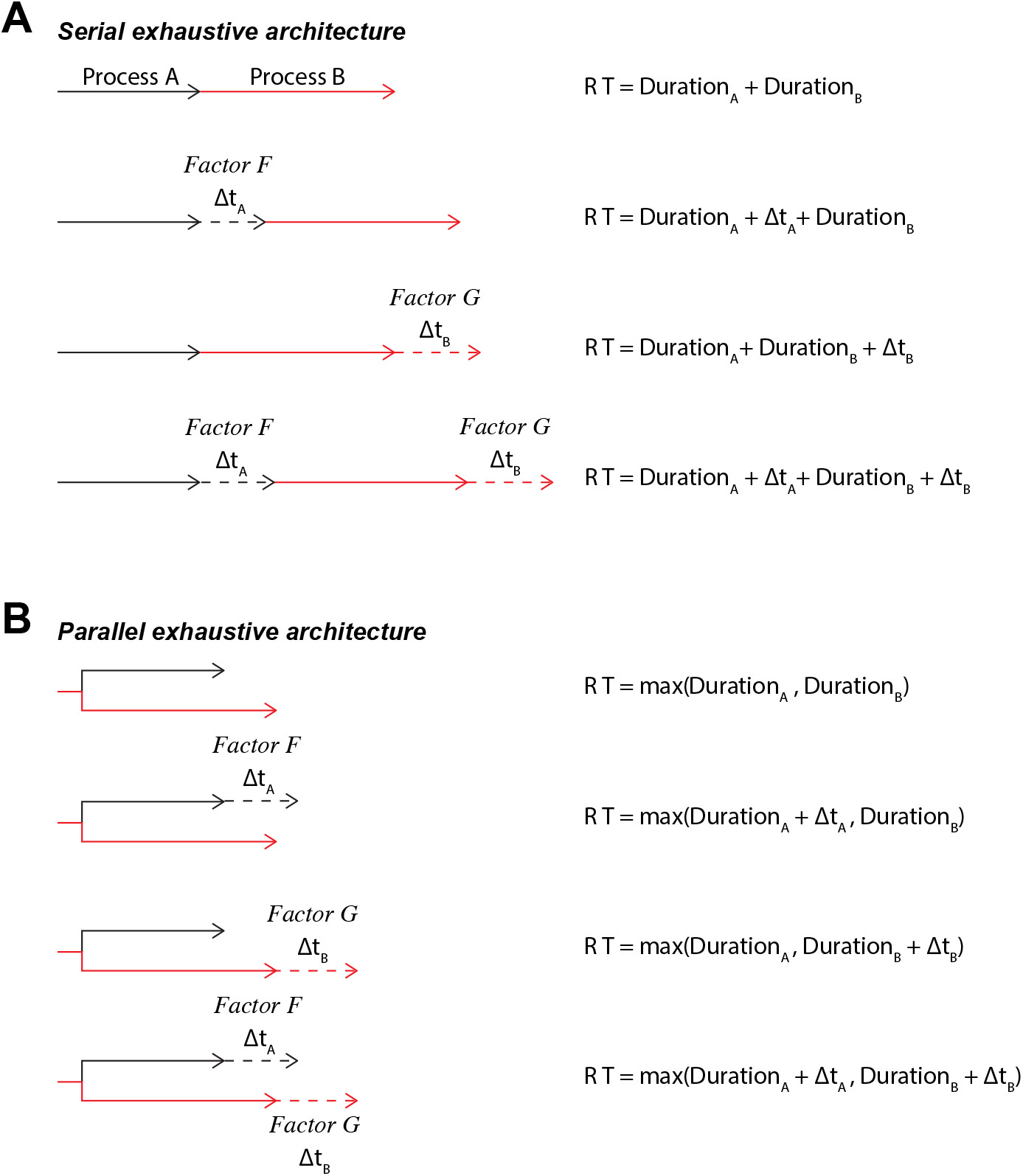
Two alternative architectures for the interaction of two distinct processes. (**A**) Serial exhaustive architecture. Both processes must complete before a response can be initiated. The durations of the two stages of processing, A and B, are under the selective influence of factors, *F* and *G*. Mutual invariance is satisfied when manipulation of factor *F* (or *G*) alters the duration of stage A (or B) but not B (or A). Additivity is satisfied when the total RT equals the sum of the durations of the separate processes. (**B**) Parallel exhaustive architecture. The two processes operate concurrently but both must complete before a response can be initiated. Manipulation of factor *F* (or *G*) alters the duration of stage A (or B) but not B (or A). The variation of RT across the two manipulations is additive or under-additive.

During cognitive neurophysiological experiments, RT can be divided into distinct processing stages during visual search (Thompson et al. 1996; cf. Costello et al. 2013). The singleton selection stage takes longer when the target is more similar to distractors (Sato et al. 2001). Saccade preparation is delayed by less efficient visual search (Woodman et al. 2008). Requiring arbitrary stimulus-response mapping reveals more functional categories of neurons because it requires more computational operations that occupy different intervals including singleton selection time, saccade endpoint selection time, and the time when the stimulus-response rule was registered (Sato and Schall 2003; Schall 2004).

Crucially, single-stage models cannot explain tasks that require multiple, sequential operations. Consider a visual search filtering task that requires a “decision” about the location of a color singleton, a “decision” about the shape of the singleton, a “decision” about the shapes of distractors, a “decision” about the congruency of the singleton and distractor shapes, a “decision” about the instructed stimulus-response mapping, a “decision” about the correct endpoint of the saccade, and a “decision” about when to initiate the saccade. Below we will describe performance of monkeys in such a task.

The literature is divided on how filtering tasks are performed. The most common view is that selecting an object and categorizing it are separate sequential stages (Figure 1A) (Broadbent 1971; Hoffman, 1978; Treisman 1988; Wolfe 2014). An alternative view is that objects are selected and categorized through parallel processes (Figure 1B) (Bundesen 1990; Logan 2002). The fundamental problem of distinguishing serial from parallel processing has proven challenging because particular serial and parallel architectures can be mathematically indistinguishable (e.g, Townsend 1972, 1990). Hanes & Schall (1996) proposed that the patterns obtained in neural recordings can resolve such model mimicry. This paper outlines an approach to elucidating processing architectures accomplishing a visual search task.

## METHODS

### Subjects

Behavioral data were collected from two macaque monkeys, *Macaca mulatta* and *M. radiata*, identified as Le and Da. The monkeys weighed approximately 12 kg (Le) and 8 kg (Da) and were aged 6 years (Le) and 12 years (Da) at the time of the study.

### Surgical procedures

Monkeys were surgically implanted with a headpost affixed to the skull via ceramic screws under aseptic conditions with isoflurane anesthesia. Antibiotics and analgesics were administered postoperatively. Monkeys were allowed at least 6 weeks to recover following surgery before being placed back on task. All procedures were approved by the Vanderbilt Institutional Animal Care and Use Committee in accordance with the United States Department of Agriculture and Public Health Service Policy on Humane Care and Use of Laboratory Animals.

### Task design and protocol

Monkeys performed 30 sessions of a go-nogo visual search task in which response was cued by the shape of a color singleton. Trials began with the monkey fixating a central stimulus for 800-1200 ms, after which eight iso-eccentric, isoluminant stimuli were presented with eccentricity = 6.0 deg. Stimuli had three possible shapes: square (aspect ratio = 1.0), or rectangle (aspect ratio = 1.4 or 2.0). All eight stimuli had the same shape on each trial. If the singleton and distractors were square (i.e. no-go trial), monkeys were rewarded for maintaining fixation at the central spot for ~ 1 sec; if stimuli were rectangular, monkeys were rewarded for shifting gaze to the singleton and maintaining fixation for 800 (monkey Le) or 1000 (monkey Da) ms.. The inter-trial interval was fixed at 2 sec.

Task difficulty varied along two dimensions (Figure 2): singleton-distractor color similarity and stimulus shape discriminability (i.e. aspect ratio). All stimuli had four possible colors: red, off-red, green, and off-green. The orientation of elongation was counterbalanced between the two monkeys; for monkey Da a vertical rectangle signaled go, whereas for monkey Le a horizontal rectangle signaled go. Task difficulty increased with high singleton-distractor color similarity and low stimulus aspect ratio. No-go trials comprised ~20% of all trials in each session.

**Figure 2.**
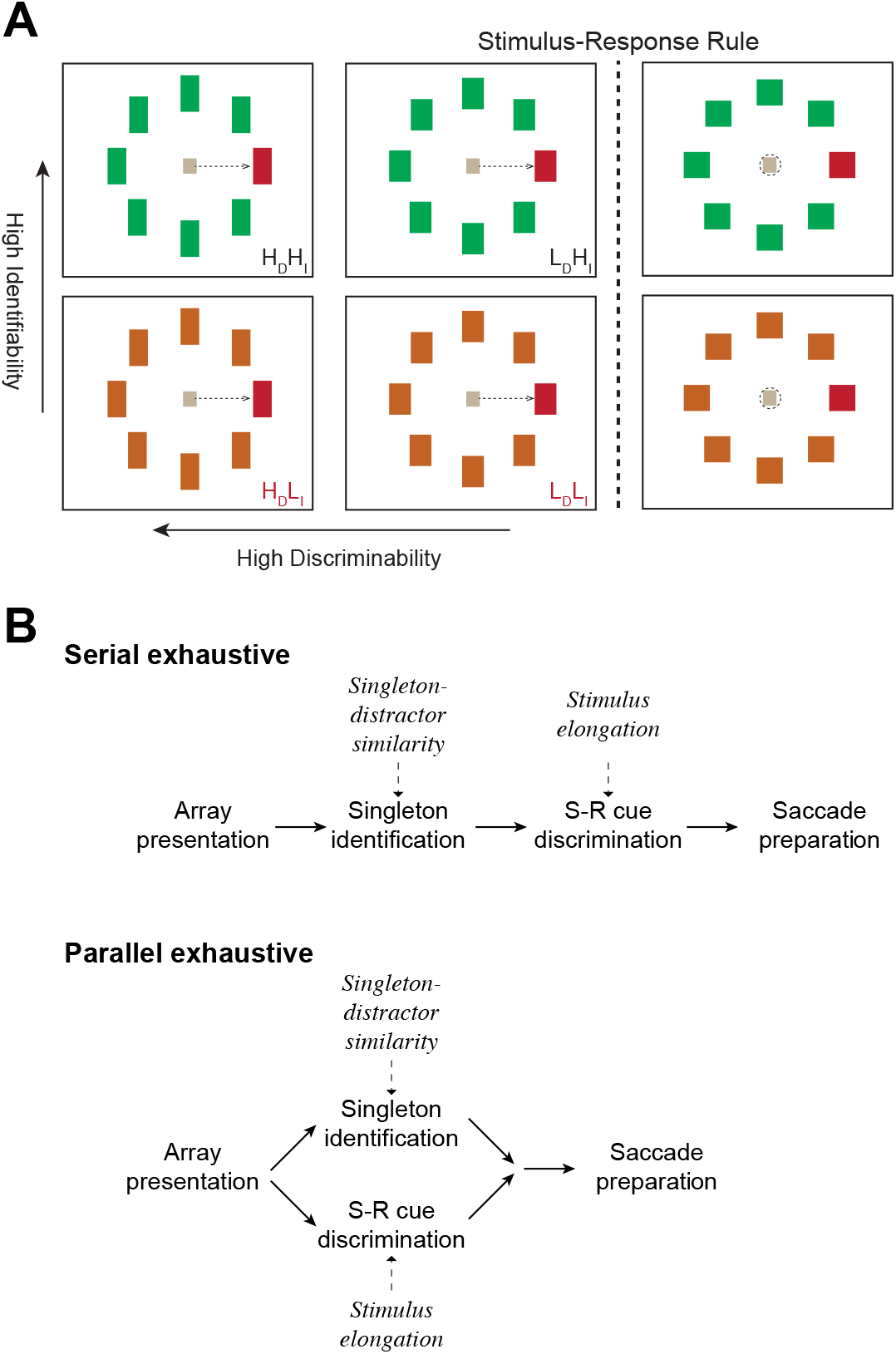
Visual search task designed to elucidate distinct operations. (**A**) Visual search task with go-nogo stimulus-response mapping. Six representative trial types are depicted. Correct gaze behavior is illustrated with dotted arrows for go trial saccades or dotted circle for nogo maintained fixation. The singleton is illustrated as always red and located on the right for purposes of illustration. Singleton shape cued the response rule. If the singleton was square (*right*), it cued withholding of the saccade. If the singleton was elongated (*left* and *middle*), it cued a pro-saccade. Two factors were manipulated independently. Stimulus-response cue discriminability was either High (aspect ratio = 2.0, H_*Discrim*_ or H_D_) or Low (aspect ratio = 1.4, L_*Discrim*_ or L_D_). On each trial, all distractors shared the degree of elongation with the color singleton. Singleton identifiability was either High (larger chromatic difference between singleton and distractors, H_*Ident*_ or H_I_) or Low (smaller chromatic difference between singleton and distractors, L_*Ident*_ or L_I_). The task offered 4 basic types of trials: High Identifiability and High Discriminability (H_*Ident*_H_*Discrim*_), Low Identifiability and High Discriminability (L_*Ident*_H_*Discrim*_), High Identifiability and Low Discriminability (H_*Ident*_L_*Discrim*_), and Low Identifiability and Low Discriminability (L_*Ident*_L_*Discrim*_). To assess the additivity and mutual invariance of these factors, trial types were interleaved in a 2×2 design. To assess how each factor affected processing capacity, trial types were also interleaved in blocks of 2×1 and 1×2 design: (H_*Discrim*_ and L_*Discrim*_ with only H_*Ident*_) and (H_*Discrim*_ and L_*Discrim*_ with only L_*Ident*_) and 1×2 interleaved trials: (H_*Ident*_ and L_*Ident*_ with only H_*Discrim*_) and (H_*Ident*_ and L_*Ident*_ with only L_*Discrim*_). (**B**) Alternative processing architectures for the double factorial visual search task. Singleton identification is influenced by target-distractor similarity but not singleton elongation. Stimulus-response cue discrimination is affected by singleton elongation, but not target-distractor similarity. Under the serial exhaustive architecture (top), singleton identification is completed before cue discrimination, which must then be completed before production of the response. Under the parallel exhaustive architecture (bottom) singleton identification and cue discrimination operate concurrently and must both finish before production of the response.

### Gaze data acquisition and analyses

We used an Eyelink 1000 system (SR Research; sampling rate□=□1,000 Hz) to track the left eye of both monkeys.

### Assessment of operations, stages and strategies

To assess quantitatively alternative process architectures supporting performance of this task, we applied systems factorial technology (Townsend & Nozawa, 1995; Harding et al. 2016), which has been employed to investigate human visual search performance (Fifić et al. 2008b). Systems factorial technology is based on mathematical propositions specifying experimental conditions offering strong tests of alternative architectures. Under conditions of selective influence, predictions of serial and parallel models with different decisional stopping rules are distinct. Through a series of particular analyses of response time distributions, systems factorial technology can discriminate between five types of information processing architectures that could accomplish a task. These are *serial self-terminating*, *serial exhaustive*, *parallel self-terminating*, *parallel exhaustive*, and *coactive*. Distinguishing serial from parallel processing in visual search has a long history (e.g., Treisman & Gelade 1980; Townsend 1990), yet progress on this issue remains possible; through systems factorial technology, when selective influence is applied effectively, then predictions of serial and parallel models and their stopping rules are mathematically distinct and experimentally discriminable. Systems factorial technology requires a 2×2 manipulation of factors that influence processing. Thus, as illustrated in Figure 2, for this study, the first manipulation was target-distractor similarity, or *singleton identifiability*, with low similarity and high similarity search arrays, which we term High Identifiability (H_*Ident*_) and Low Identifiability (L_*Ident*_) respectively. The second manipulation was *cue discriminability* of the singleton with High Discriminability (H_*Discrim*_) and Low Discriminability (L_*Discrim*_) according to the aspect ratio of the singleton. This 2×2 design results in four trial types, ranging from the easiest High Identifiability and High Discriminability (H_*Ident*_H_*Discrim*_) to the most difficult Low Identifiability and Low Discriminability (L_*Ident*_L_*Discrim*_).

### Statistical analyses

All t-tests presented are two-sided, unless otherwise stated.

## RESULTS

### Monkeys are sensitive to cue discriminability and singleton identifiability

Response times (RT) of both monkeys in this study were affected by both task manipulations. The average RT for each condition are shown in Figure 3A. As expected response times were longer for trials in which the singleton was more chromatically similar to distractors and thus easier to identify (Monkey Da: Low Identifiability (L_*Ident*_) = 272 ± 13 ms, High Identifiability (H_*Ident*_) = 236 ± 23 ms, F_(1,116)_ = 246.63, p < 0.01; Monkey Le: L_*Ident*_ = 263 ± 31 ms, H_*Ident*_ = 227 ± 31 ms, F_(1,116)_ = 161.12, p < 0.01). Likewise, average RT was longer when the cue was less discriminable (Monkey Da: Less discriminable (L_*Discrim*._) = 281 ± 23 ms, More discriminable (H_*Discrim*._) = 228 ± 14 ms, F(1,116) = 535.09, p < 0.01; Monkey Le: L_*Discrim*._ = 264 ± 34 ms, H_*Discrim*._ = 227 ± 11 ms, F(1,116) = 170.17, p < 0.01). These factors had a significant interaction in both monkeys (Monkey Da: F(1,116) = 11.25, p < 0.01; Monkey Le: F(1,116) = 6.52, p = 0.012).

**Figure 3.**
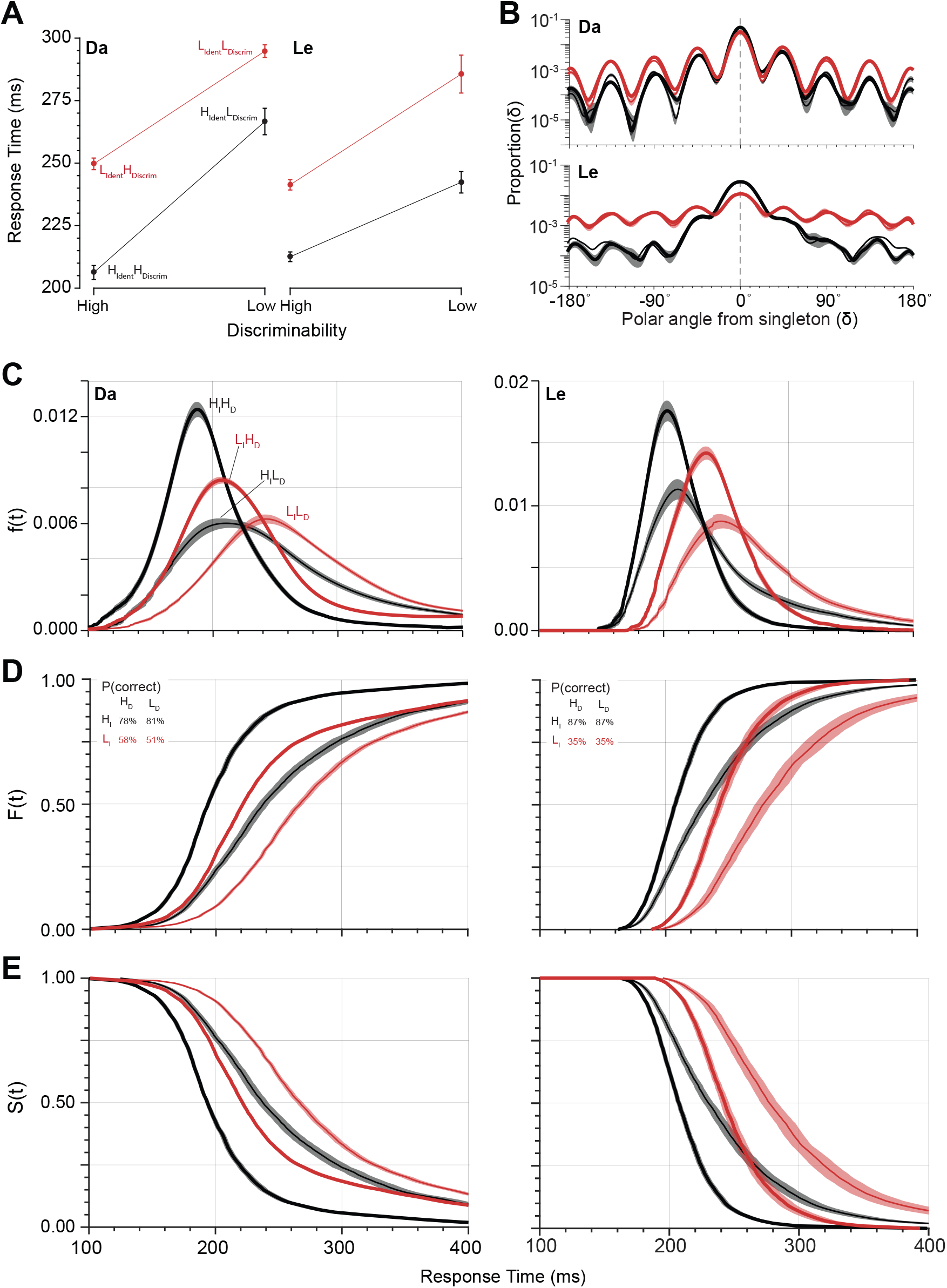
Basic performance measures. (**A**) Mean RT ± SEM for each trial type of the double factorial paradigm. There were four trial types: H_*Ident*_H_*Discrim*_, H_*Ident*_L_*Discrim*_, L_*Ident*_H_*Discrim*_, and L_*Ident*_L_*Discrim*_ Trials with High and Low singleton identifiability are shown in black and red, respectively. Monkey Da exhibited under-additivity of RT, whereas monkey Le exhibited over-additivity of RT across the two manipulations. (**B**) Log plot of the probability density of saccade endpoints relative to singleton location for High (black) and Low (red) singleton identifiability and High (bold) and Low (thin) cue discriminability. Spacing between search stimuli was 45° in polar angle. Both monkeys exhibited higher incidence of error saccades to the location adjacent to the singleton. Error bands are SE across sessions. (**C**) Probability density of RT, f(t). (**D**) Cumulative distribution of RT, F(t). Percent correct for each trial type is inset. (**E**) Survivor function of RT, S(t) = 1 – F(t).

The endpoints of correct and error saccades were informative. As observed previously (e.g., Findlay 1997), on error trials both monkeys more commonly shifted gaze to a distractor adjacent to the color singleton. Both monkeys made some saccades toward the color singleton when it was a square (Da: 11.5 ± 5.2% H_*Ident*_, 11.3 ± 3.3% L_*Ident*_; Le: 26.7 ± 15.0% H_*Ident*_, 7.2 ± 7.1% L_*Ident*_). This demonstrates that the task rule was learned and that squares and less elongated rectangles were sufficiently similar to invoke difficult cue discriminability.

Saccade endpoint was also affected more by singleton identifiability than by shape discriminability (Figure 3B). Percent correct under high identifiability was greater relative to low identifiability (Da: 79.5 ± 4.3% H_*Ident*_, 54.4 ± 10.7% L_*Ident*_; Le: 87.0 ± 5.5% H_*Ident*_, 35.3 ± 8.7% L_*Ident*_). These effects were significantly different between monkeys (Da: F(1,116) = 282.98, p < 0.001; Le: F(1,116) = 1506.05, p < 0.001). The effect of cue discriminability on saccade endpoint was different for the two monkeys. Both monkeys were equally accurate when stimulus shape was more discriminable (Le: 61.2 ± 6.4% H_*Discrim*._, 61.2 ± 8.1% L_*Discrim*._, F(1,116) = 0, p = 0.9643; Da: 64.8 ± 7.7% H_*Discrim*._, 69.1 ± 8.6% L_*Discrim*._, F(1,116) = 8.35, p < 0.01).

Singleton identifiability and cue discriminability also influenced the shape of the RT distributions. To prepare for the systems factorial analysis, we illustrate the variation of the RT distributions in probability densities (f(t) = Prob(t < RT < t+Δt) (Figure 3C), cumulative distributions (F(t) = ∫f(t)dt = Prob(RT≤t)) (Figure 3D), and survivor functions (S(t) = Prob(RT > t) = 1 – F(t)) (Figure 3E). The influence of both factors on the characteristic shape of these RT distributions is clear for both monkeys. However, much deeper computational insights are available through the next steps of analysis.

### Systems factorial technology-based assessment of visual search performance

Systems factorial technology is used to assess processing stage architecture and performance strategy by analyzing the RT distributions of each condition within a 2×2 factorial design (Townsend and Nozawa 1995; Houpt et al. 2014; Harding et al. 2016). Given that each factor (singleton identifiability and color discriminability) affected RT, we assessed the manner in which one factor affected RT while the other factor was fixed. That is, the way that shape affects RT on trials where singleton and distractors were dissimilar in color may or may not be the same as the way that shape affects RT on trials where singleton and distractors were similar in color.

Figure 4 illustrates the rationale and implementation of systems factorial technology using a system of simple simulations. The 5 alternative architectures were simulated with pairs of simple linear accumulators embodying two processes, designated **A** and **B** (Carpenter & Williams 1995; Brown & Heathcote, 2008). The finishing times of the accumulators were defined by four key parameters, the threshold, drift rate, drift rate variability, and non-decision time. For the current purposes, we simulated each processing stage by assigning an arbitrary threshold and a non-decision time of zero. An arbitrary mean drift rate was assigned for the more efficient condition of each factor, and a slower drift rate was assigned for the less efficient condition of each factor. Each manipulation was also assigned identical drift rate variability. For the combined manipulation, the drift rate effects were added. Each replicate for each condition had a drift rate sampled from a normal distribution centered on the assigned mean drift rate and with a standard deviation of the assigned drift rate variability. The parameters of each simulation were adjusted to produce similar ranges of RT. The resultant process durations were assessed by 10000 random samples defined by each manipulation’s drift rate parameterization.

**Figure 4.**
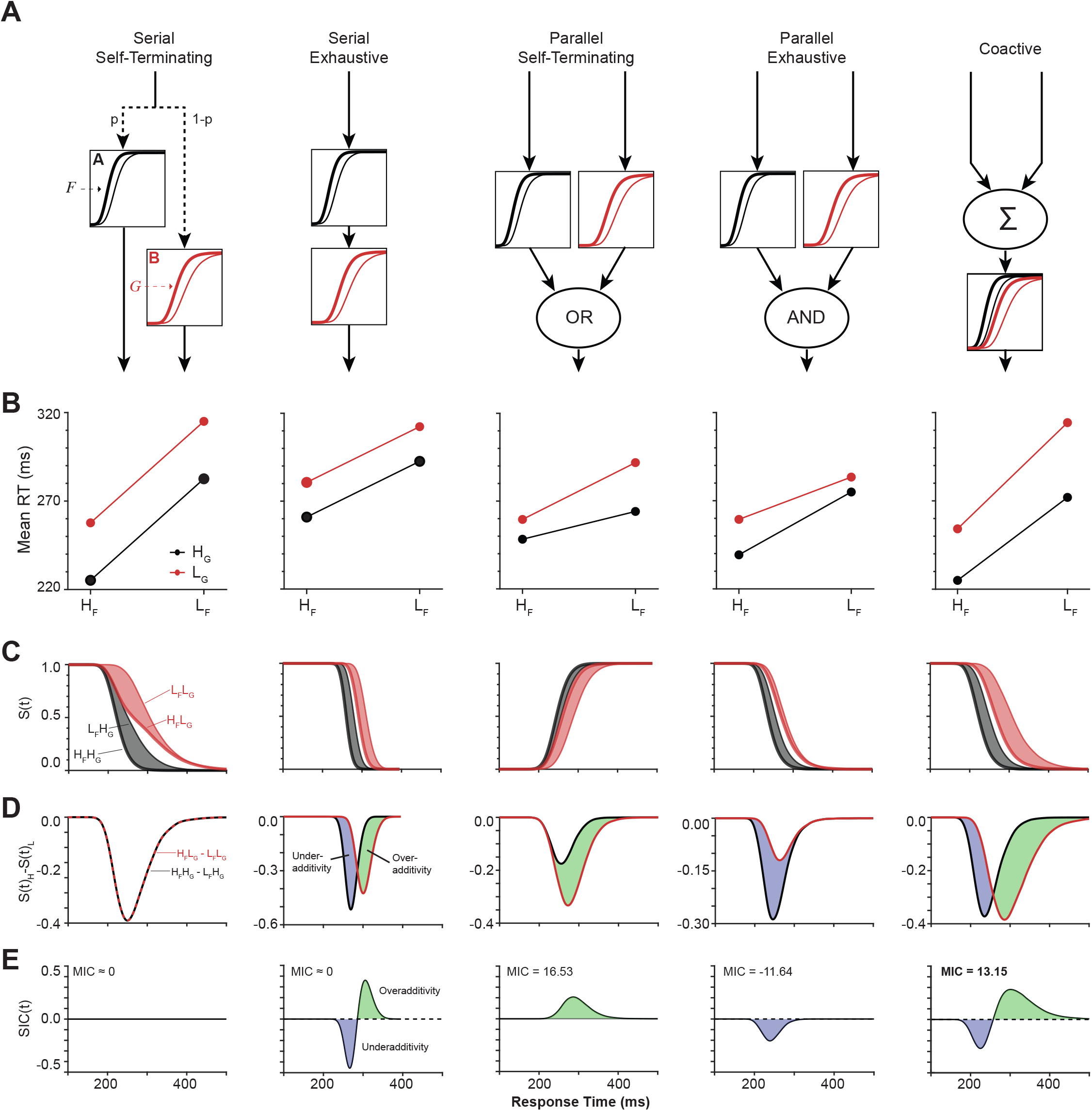
Systems factorial technology simulations. (**A**) Each of five processing architectures were modeled using two simple linear accumulator models, each representing an independent operation or stage. The two operations, A and B, were assumed to be under the selective influence of Factors *F* and *G*. Stage A varied with Factor *F* (but not *G*), and Stage B varied with Factor *G* (but not *F*). Essential features of each architecture are shown with depictions of relative stage durations. (**B**) Mean interaction contrast. Plots of mean RT for each trial type of the double factorial setup. Lines in red and black refer to Low and High levels of Factor *G*. (**C**) Survivor function *S(t)* for each trial type. The gray and red shadings highlight the effects of Factor *F* on *S(t)* at fixed levels of Factor *G*. (**D**) Difference in survivor function *S(t)* for fixed levels of Factor *G*. Regions of blue and green denote intervals of underadditivity and overadditivity, respectively. (**E**) Survivor interaction contrast *S(t)*. The serial self-terminating architecture produced a SIC that did not differ from 0.0 for all time. The serial exhaustive architecture produced a SIC that deviated to under-additivity followed by over-additivity, with equal area under each region. The parallel self-terminating architecture produced a SIC with overadditivity. The parallel exhaustive architecture produced a SIC with underadditivity. The coactive architecture produced a SIC that deviated to underadditivity followed by overadditivity, with greater area under the overadditive region for net overadditivity.

We explore the influence of two factors, designated *F* and *G*, which cause highly efficient (H) and less efficient (L) processing. For example, factor *F* could be singleton-distractor similarity that influences the duration of singleton identification (process **A**), and factor *G* could be singleton shape that influences the duration of cue discrimination (process **B**). We now present the 5 possible processing architectures resolved by SFT.

Consider first processes **A** and **B** as serial self-terminating processes (Figure 4A). The two processes are queued sequentially, but only one needs to be completed for the overt response to be produced. Formally, the order of sub-processes is unknown and random. The two levels of factor *F* result in two distributions of process finishing times that overlap but have different modal values. Similarly, the two levels of factor *G* result in two distributions of finishing times that overlap but have different modal values. In this architecture, RT on each trial corresponds to the finishing time of the fastest process. Of course, process **A** or **B** might finish first on a given trial, but on average the systematic variation of RT will depend on the influence of the respective factors on each process. Under this architecture the influence of each factor on each process is independent. This results in mutually invariant, additive differences in average RT (<RT>) of both processes across both factors. In other words, a plot of average RT produced for each combination of the 2×2 design will produce parallel relations with no interaction across factors. The nature of the interaction across factors can be summarized by a value known as the *Mean Interaction Contrast* (MIC), which is calculated as

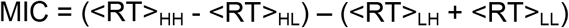

where <RT>_HH_ is the mean RT on trials with both factors allowing high efficiency for their respective processes, <RT>_LL_ is the mean RT on trials with both factors allowing only low efficiency for their respective processes, and <RT>_HL_ and <RT>_LH_ are the mean RT on trials with one factor allowing high efficiency for its process with the other factor allowing only low efficiency for its process.

For the <u>serial self-terminating</u> processes, MIC = 0, which indicates perfect additivity of the underlying processes. Non-zero values of MIC signify an interaction among the processes. Such an interaction can be underadditive (MIC < 0) or overadditive (MIC > 0). MIC > 0 identifies either parallel self-terminating or coactive process architectures, and MIC < 0 identifies parallel exhaustive processes. Thus, the MIC offers some insight into the nature of the interaction between sub-processes. However, MIC cannot discriminate between the self-terminating or exhaustive stopping rules for serial architectures or discriminate between coactive and parallel self-terminating architectures (Townsend and Nozawa 1995).

Further insight is available through examination of the entire distribution of RTs across conditions. The effects of the combination of conditions can be assessed as a function of time over the production of the responses by measuring the difference of the survivor functions. Given the independence of the influences of the 2 factors, the rate of response production through time for each process does not vary across each level of the other. This can be quantified by measuring the effect of one factor within each condition of the other factor; the difference between response production when one factor is highly efficient (H_F_) and when it is less efficient (L_F_), while the other factor is more (G_H_), or less (G_L_), efficient. For example, the effect of varying factor *F* with respect to varying factor *G* can be assessed by subtracting S(t) for *H*_*F*_ and *L*_*G*_ from S(t) for *H*_*F*_ and *H*_*G*_ and also subtracting S(t) for *L*_*F*_ and *L*_*G*_ from S(t) for *L*_*F*_ and *H*_*G*_. The interaction between the two manipulations, known as the *survivor interaction contrast (SIC).* The SIC is a distribution-free measure for assessing the architecture (i.e., serial or parallel) and stopping rule (i.e., minimum time or maximum time) of information processing, which indexes the difference in levels of *G* between the levels of *F*, is calculated similar to the MIC by subtracting the two resulting difference functions over time:

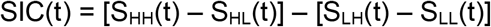

where the value SIC(t) at a given time t, S_HH_(t) is the value of the survivor function at time t when both factors are more efficient (H_F_H_G_), S_LL_(t) is the value of the survivor function at time t when both factors are less efficient (L_F_L_G_), S_HL_(t) is the value of the survivor function at time t when factor F is more efficient and factor G is less efficient (H_F_L_G_), and S_LH_(t) is the value of the survivor function at time t when factor F is less efficient and factor G is more efficient (L_F_H_G_). These operations are commutative thus the effect of varying *G* with respect to varying *F* is expected to be equivalent. The SIC measures the interaction contrast throughout the duration of all processes. The basic concepts of additivity, underadditivity, and overadditivity apply to the SIC; they just apply through time. Under the assumptions of systems factorial technology (e.g., stochastic independence of the processes), the form of SIC(t) is diagnostic of the 5 processing architectures.

The purely additive influence of factors in the serial self-terminating architecture result in SIC values that do not vary over time. However, the SIC produced by the other 4 architectures varies through time, each producing a different pattern of variation. Accordingly, the pattern of variation of the SIC curves can diagnose which underlying architecture produced a given pattern of RTs in the 2×2 experimental design.

Consider the serial exhaustive architecture. The processes are queued sequentially and the overt response is produced only when both processes have finished. Formally, the order of processes is unknown and SFT is unable to identify which one acted first. The mean RTs across factors exhibit no sign of interaction, so the MIC = 0 for this architecture as well. However, through time this architecture produces first underadditivity then overadditivity. That is, the SIC exhibits a negative-going followed by a positive-going deflection. Importantly, to satisfy the requirement that MIC = 0, the areas under the negative-going and positive-going deflections are equivalent.

Consider next the parallel self-terminating architecture. Both processes operate simultaneously, so a stopping rule must be specified. Specifically, if a response can be made when one stage is complete, then the combined process is parallel self-terminating. In this architecture, the overt response is produced as soon as either process finishes. This architecture is also known as a race and predicts overadditivity. Thus, MIC > 0, and the SIC curve deviates only positively.

Next, consider the parallel exhaustive architecture in which a response can only be made when both stages are complete. Both processes operate simultaneously, but the overt response is produced only after both processes have finished. This architecture predicts underadditivity. Thus, MIC < 0, and the SIC curve deviates only negatively. The performance of one of the monkeys will have this appearance.

Consider, finally, the coactive architecture. While more complex and less explicit in form, it can be distinguished in function through these methods. The core idea is that the two processes interact in a manner that can be characterized as finer grain coordination such as summation of the respective states through time. This can be realized if neither of the two processes **A** nor **B** produce the overt response but instead provide activations to a third process that sums the activations from **A** and **B** and hence produces the overt response. This architecture, like a serial exhaustive architecture, predicts first underadditivity and then overadditivity. However, unlike a serial exhaustive architecture, for the co-active architecture, MIC > 0. Therefore, the area under the positive-going deflection is greater than the area under the negative-going deflection and the architecture predicts a net overadditivity. Accordingly, although this architecture has an initial negative dip and a positive deflection (like serial processing) and has an MIC greater than 0 (like parallel self-terminating), the combination of SIC and MIC differentiates it from either of these other architectures. The performance of the second monkey will have this appearance.

### Processing architectures supporting visual search

We applied the systems factorial technology to the visual search data obtained from two macaque monkeys. Figure 5A presents mean survivor functions for each level of the 2×2 factorial design for each monkey. At a fixed level of singleton identifiability, the difference between survivor functions represents the effect of shape discriminability. Figure 5B plots the difference in survivor functions for each level of singleton identifiability. The shape of these differences reveals the effect of the separate factors on response production through time. Figure 5C plots the difference of these differences, which is the survivor interaction contrast (SIC). The SIC summarizes the influence of the two factors through time.

**Figure 5.**
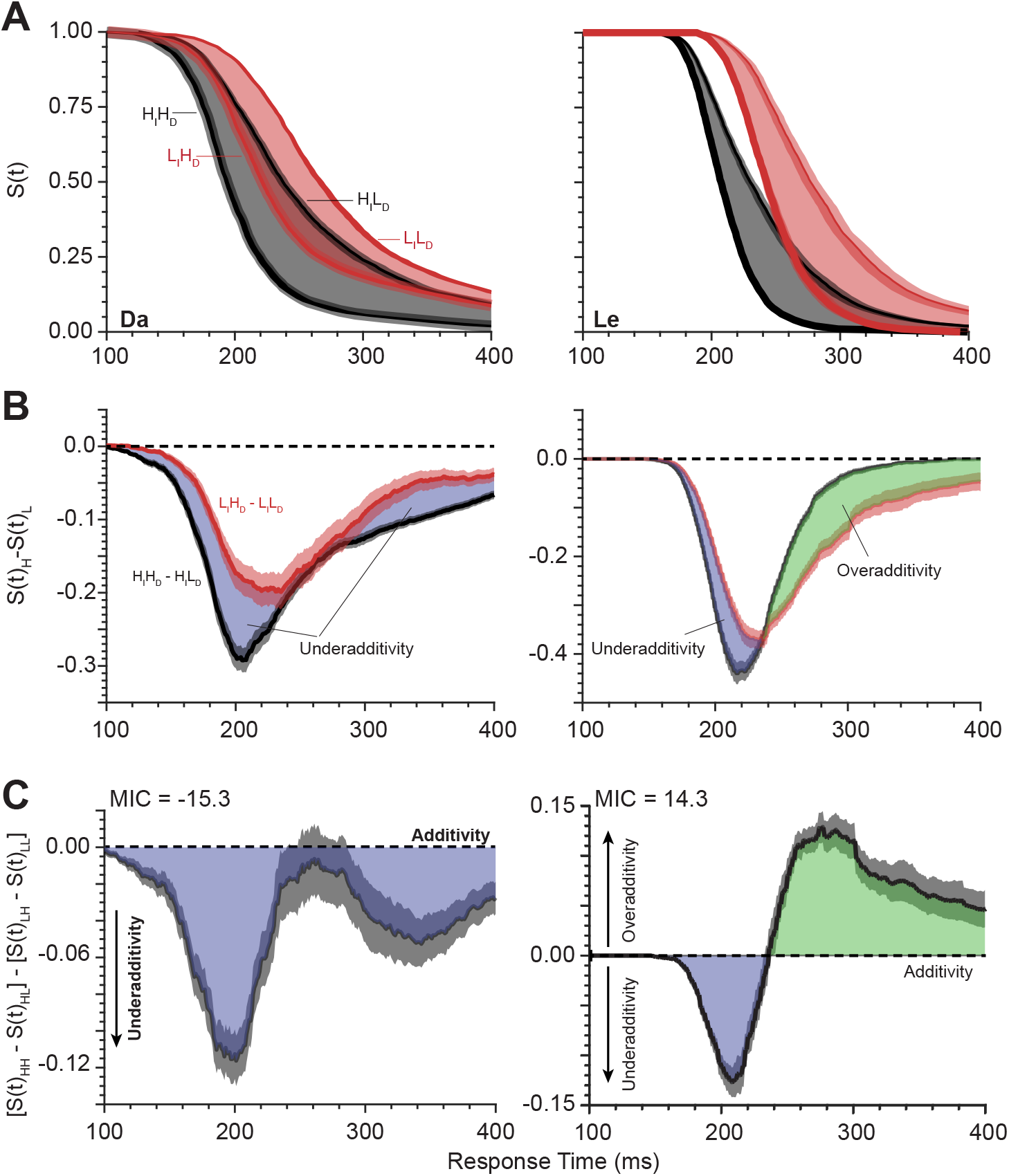
Systems factorial analysis of RT distributions from the double factorial visual search task. (**A**) Survivor functions *S(t)* for each combination of singleton identifiability and cue discriminability. Black and red lines depict High and Low singleton identifiability. Thick and thin lines depict High and Low cue discriminability. The difference between survivor functions for High and Low cue discriminability is shaded in black (High singleton identifiability) and red (Low singleton identifiability). (**B**) Difference between survivor functions for High and Low cue discriminability, computed at fixed levels of singleton identifiability. Shaded regions represent period of underadditivity (blue) and overadditivity (green) for Low (red) and High (black) singleton identifiability. Error regions are SE across sessions. (**C**) Survivor interaction contrast curves. The SIC curve for monkey Da was exclusively sub-additive, consistent with the parallel exhaustive architecture. The SIC curve for monkey Le exhibited a change from under- to over-additivity, consistent with the parallel coactive architecture.

For monkey Le, the SIC exhibited a pronounced period of underadditivity followed by a prolonged period of overadditivity. The integral of the period of overadditivity exceeded that of the underadditivity, indicative of a positive mean interaction contrast (MIC = 14.3). This outcome is characteristic of the coactive processing architecture. For monkey Da, the SIC exhibited only a prolonged and multiphasic underadditive deflection with MIC = −15.3. This outcome is characteristic of the parallel exhaustive architecture.

SFT analyses are typically performed on a per-subject basis rather than the present repeated testing across many sessions that can be done with monkeys. Thus, it is possible that the performance strategy associated with different processing architectures or dynamics varies across sessions. If so, then the multiphasic SIC curve could be an artifact of averaging sessions performed with different strategies. To assess whether the average SIC curve is a mixture of multiple architectures across different sessions, we performed a cluster analysis of SIC curves. For monkey Da, using Euclidean distance as a similarity metric identified four clusters (Figure 6A). Using correlation distance as a similarity metric was less discriminating, which indicates that the major differences in SIC are in magnitude rather than shape. To examine the systematic variability across sessions, we plotted the SIC for each cluster (Figure 6B). With MIC > 0 and SIC with a later overadditive deflection exceeding the early underadditive deflection, two of the clusters identified the coactive architecture. With MIC < 0 and only underadditive SIC deflections, the other two clusters identified the parallel exhaustive architecture. Notably, the biphasic SIC was evident in individual clusters. Even the most clearly underadditive SIC cluster had bimodal characteristics. For monkey Le, neither Euclidean nor correlation distance yielded distinct clusters. The coactive architecture was identified by the MIC and SIC values from each session, although variation in MIC magnitude varied across sessions.

**Figure 6.**
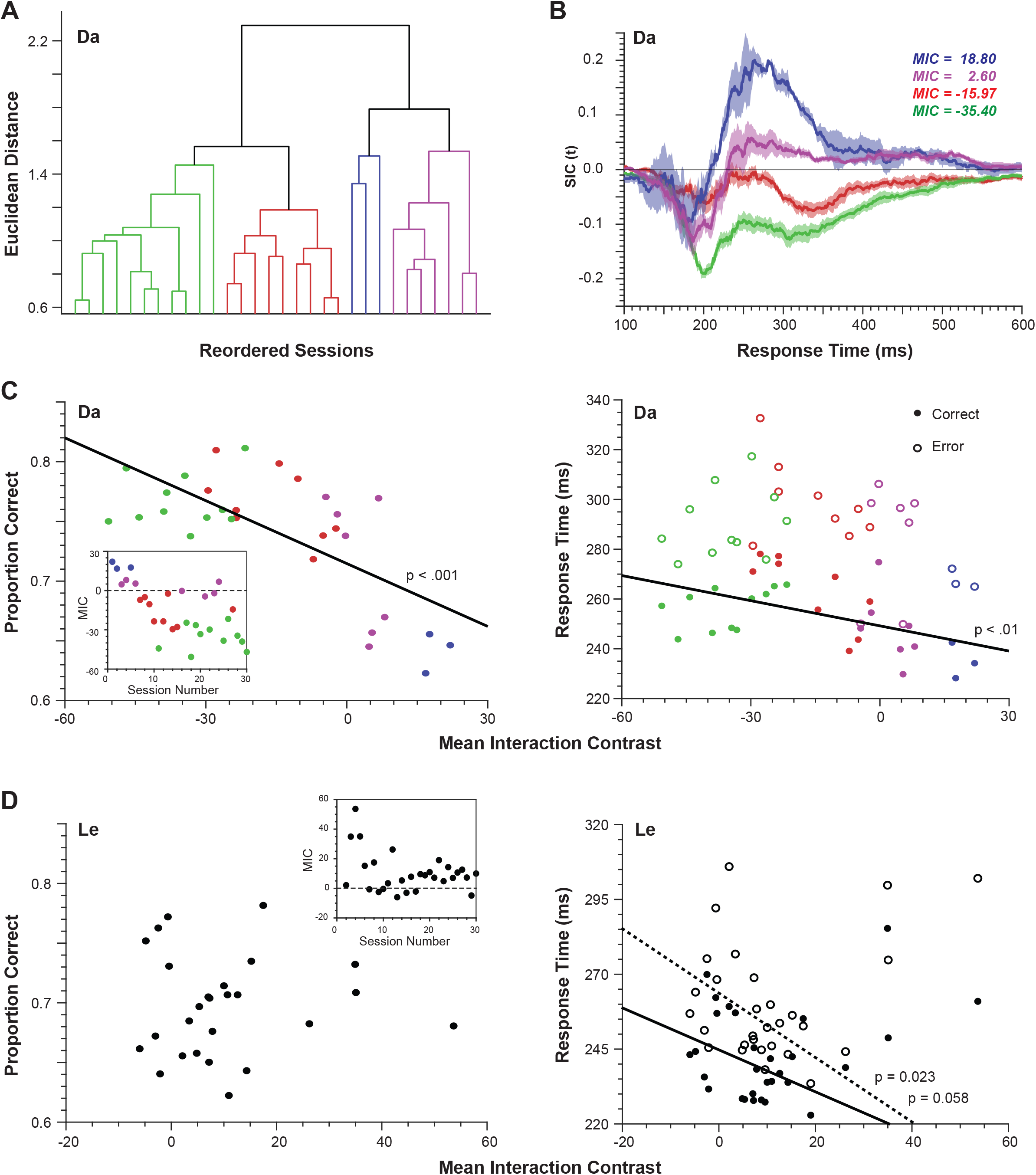
Variation of performance across sessions. (**A**) Dendrogram resulting from clustering of SIC curves across sessions for monkey Da based on Euclidean distance. Four clusters were evident, suggesting the use of different strategies. (**B**) Form of the four clusters of SIC curves. Two (blue, magenta) corresponded to the co-active architecture, and two (red, green) were unlike the SIC of any architecture. (**C**) (*left*) Proportion correct as a function of MIC across sessions for monkey Da. A strong correlation was observed. MIC across session number inset. Points are colored in accordance with their cluster identity. (*right*) Mean RT for correct (filled) and error (open) trials as a function of MIC across sessions. Error RT were longer than correct RT, and a strong correlation with MIC was observed for correct but not error RT. (**D**) (*left*) Proportion correct as a function of MIC across sessions for monkey Le. MIC across session number inset. A significant correlation was not observed. (*right*) Mean RT for correct (filled) and error (open) trials as a function of MIC across sessions. Error RT were longer than correct RT, and a correlation with MIC was observed for error RT and a trend toward correlation with MIC was observed for correct RT.

The variation in MIC values across sessions offers a unique opportunity to assess whether qualitative or quantitiative differences in processing strategies result in predictable differences in performance. Hence, we examined the relationship between the per-session MIC and the accuracy and response times during those sessions (Figure 6C). For monkey Da, we found significant negative correlations between percent correct and MIC (r = −0.69, p < 0.001) as well as between RT of correct responses and MIC (r = −0.49, p < 0.01). We found no relationship between MIC and RT on error trials (r = −0.30, p = 0.11).

For monkey Le, some early sessions had MICs clearly much greater than the majority of sessions. After considering these as outliers, we found no relationship between percent correct and MIC (r = −0.21, p = 0.32), but RT and MIC trended toward a significant negative correlation (r = −0.38, p = 0.058) for correct RTs and were significantly negatively correlated (r = −0.45, p = - 0.022) for error trials.

### Processing architectures for correct and error performance

SFT analyses commonly assume a low error rate. The performance shown here is far from perfect. However, other investigators have demonstrated that conclusions from SFT are reliable in spite of error rates approximating what we obtained (Fifić et al. 2008a). Finding the relationship between search performance and MIC values across sessions, we assessed whether performance strategies differed between correct trials and errors. Further, given the prevalence of erroneous saccades to the distractor adjacent to the singleton, we distinguished two categories of errors: informed errors made to the stimulus adjacent to the singleton and guess errors made to any other location.

Figure 7 illustrates the progression of distributions used for the SFT analysis for correct responses, informed errors, and guesses for both monkeys. The factorial manipulation trial types were assigned according to the configuration of the search array; that is, H_*Ident*_, L_*Ident*_, H_*Discrim*_, and L_*Discrim*_ were assigned with respect to the identifiability and discriminability of the singleton.

**Figure 7.**
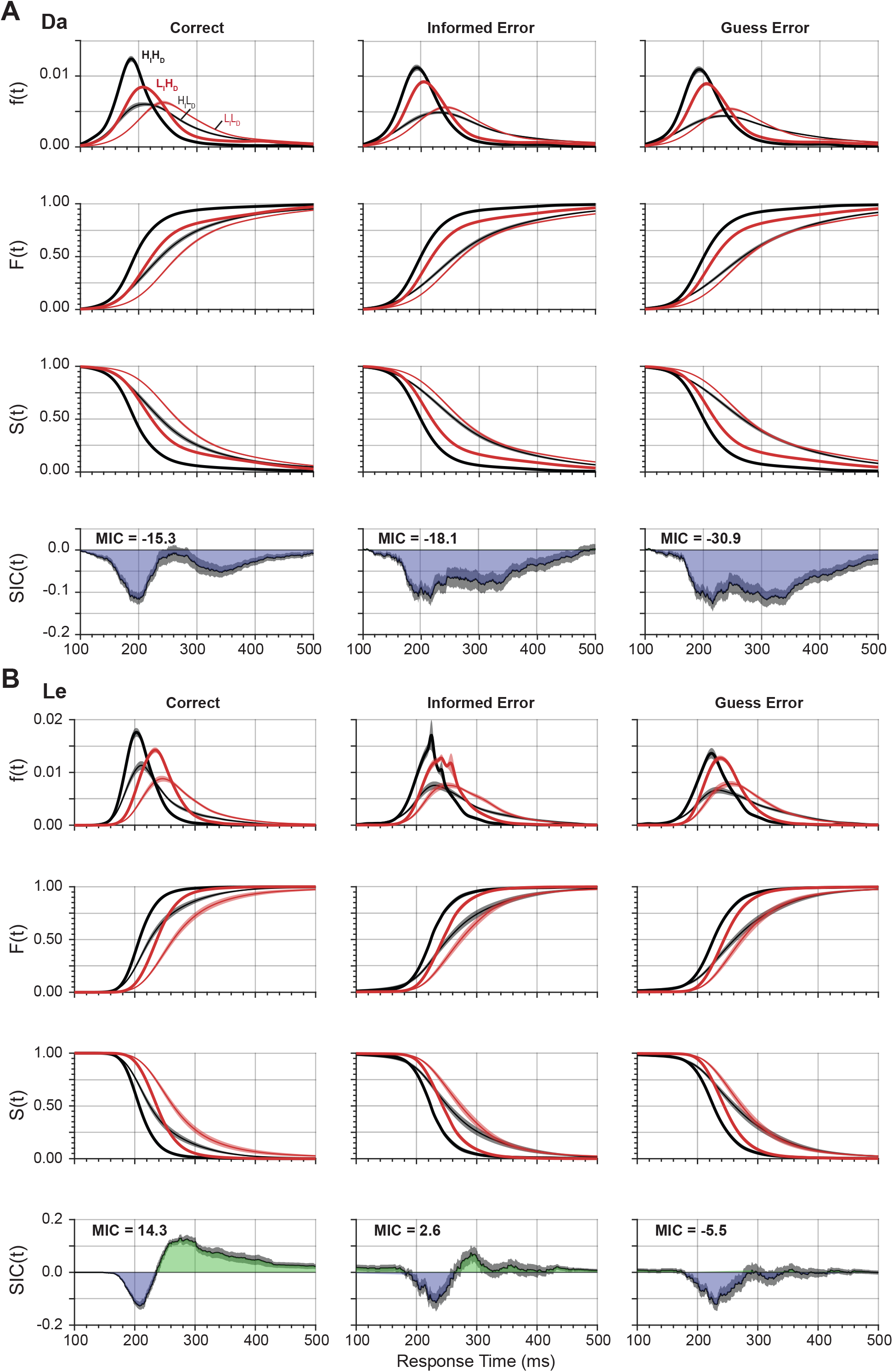
SFT analysis for different trial outcomes. (**A**) SIC curves derived from monkey Da performance on correct (left), informed errors adjacent to the singleton (middle) and guess errors (right). SIC curves for informed and guess errors exhibit more underadditivity relative to correct trials. All three SIC curves resemble that of the parallel exhaustive architecture. (**B**) SIC curves derived from monkey Le performance on correct (left), informed errors adjacent to the singleton (middle) and guess errors (right). SIC curves for errors exhibit more underadditivity (less overadditivity) relative to correct trials. The SIC curve for guess errors resembles the parallel exhaustive architecture, for informed errors, the serial exhaustive architecture, and for correct trials, the coactive architecture.

For monkey Da, both informed errors and guesses were generated with MIC < 0 and SIC deflecting only in the underadditive direction, like the correct responses. Hence, like correct responses, errors were identified with the parallel exhaustive architecture. In other words, qualitatively a single architecture produced both correct and error responses. However, quantitatively, MIC for guesses was more underadditive than MIC for informed errors, which was more underadditive than MIC for correct responses. Also, the SIC for error responses was prolonged but lacked the pronounced multiphasic pattern obtained from correct trials.

For monkey Le, we observed qualitative variation in MIC and SIC for error relative to correct trials. As noted, correct trial performance produced MIC and SIC values that identified the coactive architecture. However, for guess errors, MIC < 0, and the SIC deviated only in the underadditive direction, which identify the parallel exhaustive architecture. Meanwhile, for the informed errors, the MIC was only slightly greater than 0, and the SIC deflected more in the under-than overadditive direction. This pattern seems to approximate at least the parallel exhaustive architecture. Thus, for monkey Le, errors may originate from a processing architecture different from correct trials.

For both monkeys, although their overall SIC curves have different shapes, the MIC for correct responses (Da: MIC = −15.3; Le: MIC = 14.3) was more positive than the MIC for informed errors (Da: MIC = −18.1; Le: MIC = 2.6), which was more positive than the MIC for guess errors (Da: MIC = −30.8; Le: MIC = −5.5). It should be noted that for both monkeys this difference appears most pronounced around the time of the second negative peak in Da’s biphasic SIC curve.

### Capacity measures

The measures of SFT described above have been supplemented by measures of capacity, which quantify whether and how much the addition of a second process affects the processing efficiency of a first process. Interpretations that are ambiguous based on MIC and SIC values can be clarified using capacity measure (Eidels et al. 2011; Houpt et al. 2014; Harding et al. 2016).

Measuring capacity requires experimental conditions in which one factor is manipulated while the other is fixed. The baseline of capacity is that produced by multiple processes with unlimited-capacity, independent, and parallel. The possible outcomes of the analysis identify capacity as limited, unlimited, or super-capacity. Limited capacity processing is identified when performance improvement with increased efficiency of both processes is less than the sum of performance improvements from increased efficiency of each process independently. Unlimited capacity is identified when performance improvement with increased efficiency of both processes is equal to the sum of performance improvements from increased efficiency of each process independently. Super capacity is identified when performance improvements with increased efficiency of both processes is greater than the sum of performance improvements from increased efficiency of each process independently.

We tested both monkeys in additional sessions during which trial types were presented pseudo-randomly in a block-wise manner. We tested blocks of trials in which the 2×2 manipulations were interleaved (trial types H_*Ident*_H_*Discrim*._, H_*Ident*_L_*Discrim*._, L_*Ident*_H_*Discrim*._, and L_*Ident*_L_*Discrim*._) or each combination of 1×2 manipulations were interleaved (trial types H_*Ident*_H_*Discrim*._ and H_*Ident*_L_*Discrim*._; H_*Ident*_H_*Discrim*._ and L_*Ident*_H_*Discrim*._; H_*Ident*_L_*Discrim*._ and L_*Ident*_L_*Discrim*._; L_*Ident*_H_*Discrim*._ and L_*Ident*_L_*Discrim*._).

Figure 8 compares the various SFT performance measures for the 2×2 blocks of trials interleaved with the 2×1 and 1×2 blocks. The SIC curves derived from the trials during the 2×2 blocks replicate the patterns observed previously for both monkeys, with performance by Da identified with parallel exhaustive architecture, and performance by Le identified with co-active architecture.

**Figure 8.**
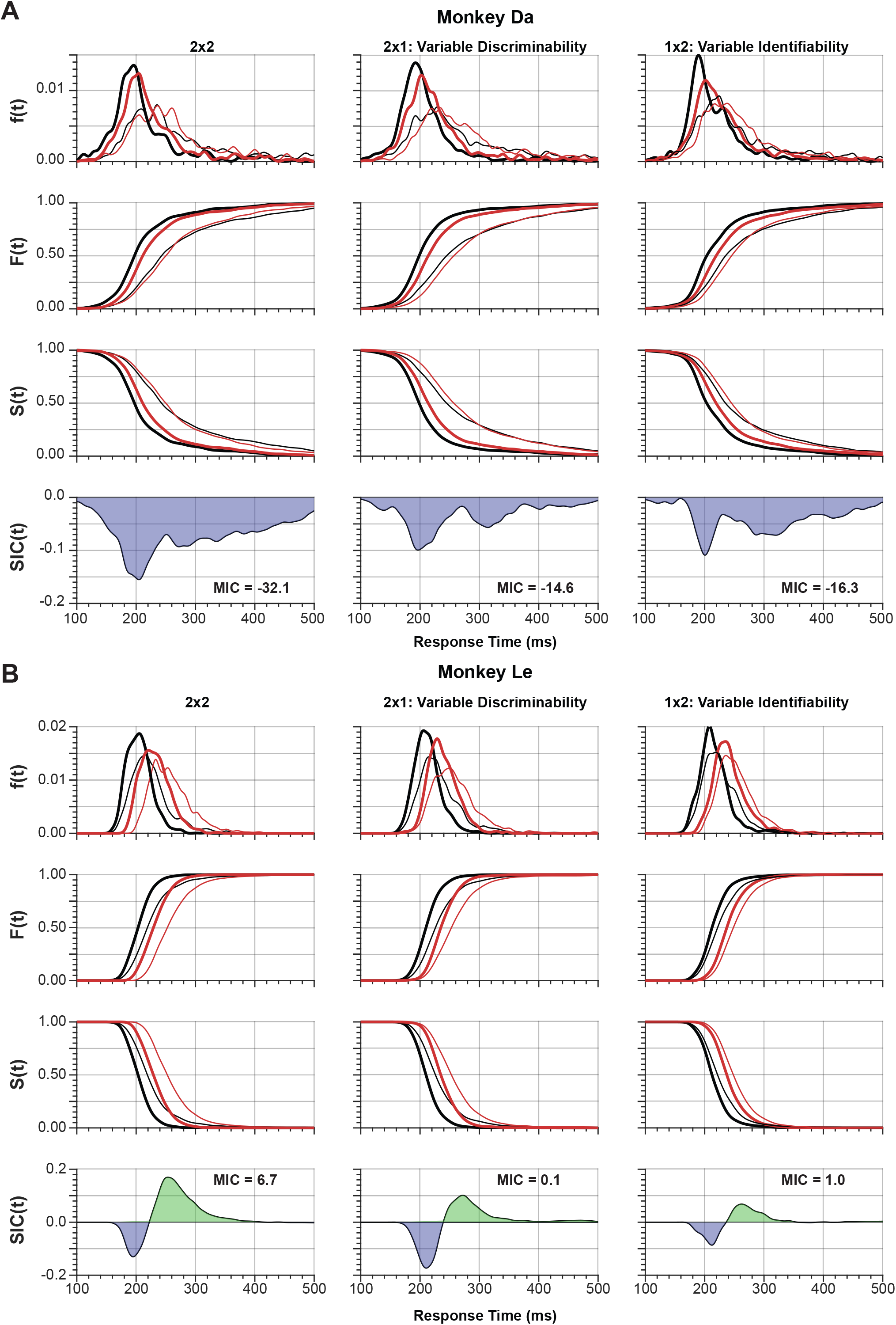
SFT analysis for single factorial paradigms. (**A**) Data for monkey Da. SIC curves were calculated for blocks of trials in which both factors could vary (2×2; *left*), blocks of trials in which only cue discriminability could vary while singleton identifiability was held constant (2×1; *middle*), and blocks of trials in which only singleton identifiability could vary while cue discriminability was held constant (1×2; *right*). The SIC curve for the double factorial paradigm (2×2) had a higher degree of underadditivity than the SIC curves for the two single factorial paradigms (2×1, 1×2). All SIC curves exhibited features resembling the exhaustive parallel architecture. (**B**) Data for monkey Le. Conventions as in (A). The SIC curve for the double factorial paradigm resembled that of a coactive architecture, whereas those of the two single factorial paradigms resembled that of a serial exhaustive architecture.

We also calculated SIC from the blocks in which identifiability was held constant (blocks of H_*Ident*_H_*Discrim*_ and H_*Ident*_L_*Discrim*_, and blocks of L_*Ident*_H_*Discrim*_ and L_*Ident*_L_*Discrim*_), and the blocks in which cue discriminability was held constant (blocks of H_*Ident*_H_*Discrim*_ and L_*Ident*_H_*Discrim*_, and blocks of H_*Ident*_L_*Discrim*_ and L_*Ident*_L_*Discrim*_). The resulting SIC curves for monkey Da also exhibited prolonged underadditive deflections, identified with parallel exhaustive processing. The smaller values of MIC in the 2×1 and 1×2 cases as compared to the 2×2 suggest that the interaction between singleton identifiability and cue discriminability is less pronounced when one factor is held constant. This appears differently in the two cases, though. In the 2×1 case where singleton identifiability is held constant and cue discriminability is variable within block, the difference between H_*Ident*_L_*Discrim*_ and L_*Ident*_H_*Discrim*_ is slightly greater than the same difference in the 2×2 blocks. In the 1×2 case where cue discriminability is held constant, the difference between H_*Discrim*_ and L_*Discrim*_ is smaller than in the 2×2 blocks, at both levels of singleton identifiability. This suggests that Da is sensitive to the block context and that eliminating one stimulus dimension from consideration reduces the interaction between the two dimensions. It also suggests that the two dimensions are treated differently in isolation; levels of singleton identifiability are more similar when interleaved whereas levels of cue discriminability are more similar when separated.

The 2×1/1×2 SIC curves for monkey Le appeared qualitatively different from that derived from the 2×2 trials. The magnitude of underadditive and overadditive deflections were nearly balanced, and the MIC values were close to 0.0. These would identify the serial exhaustive architecture (though such a transition from coactive to serial exhaustive architecture has also been found to be the result of high error rates in a coactive architecture (Fifić et al., 2008a). The differences in SIC for Le appear due to the reduction of differences between factors that are held constant. Specifically, in the 2×1 case where singleton identifiability is held constant, the difference between H_*Ident*_H_*Discrim*_ and L_*Ident*_H_*Discrim*_ is smaller than in the 2×2 case, and in the 1×2 case where cue discriminability is held constant the difference between H_*Discrim*_ and L_*Discrim*_ is smaller than in the 2×2 blocks, at both levels of singleton identifiability. Similar to Da, this suggests that Le is sensitive to the block context and that eliminating one stimulus dimension from consideration reduces the interaction between the two dimensions. Contrary to Da, this suggests that the effect of isolating one dimension or the other is the same regardless of which dimension is isolated.

Capacity values are computed using different formulae for self-terminating (C_OR_) and for exhaustive (C_AND_) architectures. The C_OR_ measure is calculated from the cumulative hazard of RT values (H(t)), which is the cumulative of minus the Log of the survivor function:

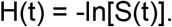

It is calculated by taking the ratio of the cumulative hazard for trials where both factors are highly efficient (H_*Ident* × *Discrim*_(t)) to the sum of the cumulative hazard for trials where each factor is highly efficient alone (H_*Ident*_(t) and H_*Discrim*_(t))

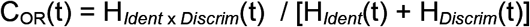

We restricted the H_*Ident*_H_*Discrim*_ trials to the 2×2 blocks of trials because the level of neither factor can be known at the start of the trial and thus must both be processed. We restricted H_*Ident*_L_*Discrim*_ trials to those in the 1×2 blocks with L_*Discrim*_ because the level of cue discriminability can be known before the trial starts and is thus removed from processing. Similarly, we restricted L_*Ident*_H_*Discrim*_ trials to those in the 2×1 blocks with L_*Ident*_ To avoid numerical complexity, we used another formulation of C_OR_(t) (Houpt et al. 2014):

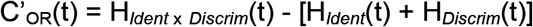

Based on the assumption of unlimited-capacity, independent, and parallel processes, with this measure, a value of 0 identifies unlimited capacity, values < 0 identifies limited capacity, and values > 0 identifies supercapacity.

The C_AND_ measure is calculated from the reverse cumulative hazard function of RT values (K(t)), which is: K(t) = log[F(t)]. C_AND_ is the ratio of the performance measured when both processes are engaged (2×2) relative to that predicted by the assumption of unlimited-capacity, independent, and parallel processes:

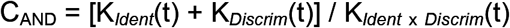

We used the alternative:

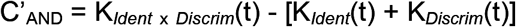

Three outcomes are possible: C_AND_ = 0 identifies unlimited capacity, C_AND_ < 0 identifies limited capacity, and C_AND_ > 0 identifies super-capacity.

Figure 9 plots the capacity measures calculated for each monkey. For monkey Da, C’_AND_ was calculated because the SIC indicated parallel exhaustive processing, and C_AND_ is used for exhaustive stopping rules. C’_AND_(t) is above 0 at all timepoints, indicating super-capacity and thus facilitation among the two sub-processes of singleton identification and cue discrimination. This is consistent with the underadditivity of the RTs when both factors are in the low condition.

**Figure 9.**
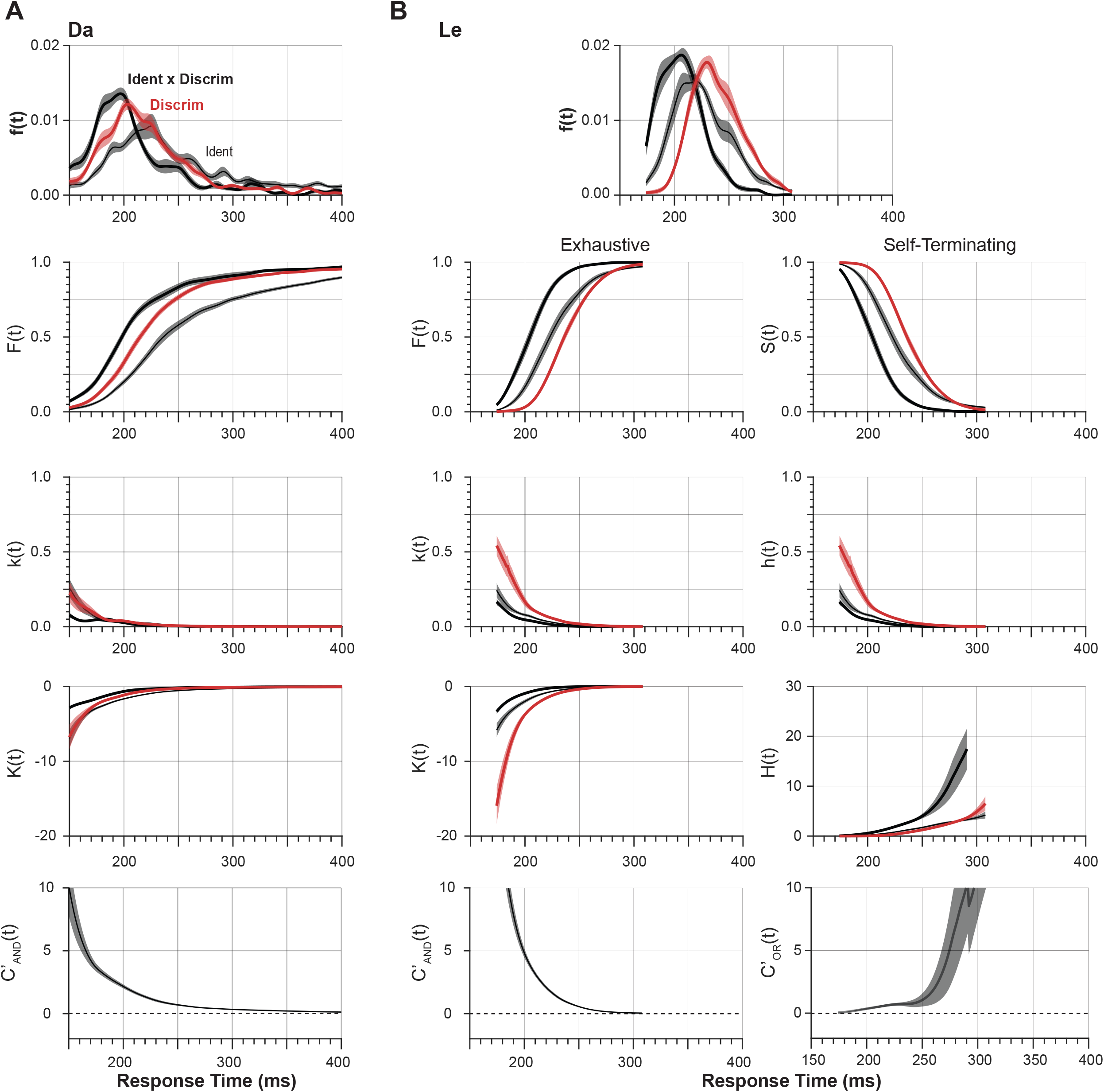
Capacity measures. (**A**) Capacity for monkey Da. Probability density functions f(RT) for the three conditions used to calculate capacity: (1) f_*Ident*_ × _*Discrim*_(RT) in which both singleton identifiability and cue discriminability were in the High condition (H_I_H_D_) in the 2×2 blocks where both factors were manipulated factorially (thick black), (2) f_*Ident*_(RT) in which singleton identifiability was High and cue discriminability was Low in 1×2 blocks where cue discriminability was invariant (H_I_L_D_), and (3) f_*Discrim*_(RT) in which cue discriminability was High and singleton identifiability was Low in 2×1 blocks where singleton discriminability was invariant (L_I_H_D_). Because the SIC curves of Da performance indicated a parallel exhaustive architecture, capacity was calculated with C’_AND_(t) based on the plotted reverse hazard function k(t), and reverse cumulative hazard function K(t). The C’_AND_ values exceeding 0 indicate super-capacity. **(B)** Capacity for monkey Le. Performance of monkey Le demonstrated a coactive architecture, thus both capacity metrics must be used. If coactive processing is exhaustive, C’_AND_(t) should be calculated (left). If, on the other hand, coactive processing is self-terminating, C’_OR_(t) should be calculated (right) based on the plotted survivor functions S(t), hazard function h(t), and cumulative hazard function H(t). Both capacity measures tended to exceed 0, indicating super-capacity.

For monkey Le, both C’_AND_ and C’_OR_ were calculated because the stopping rule of coactive processing is less clear. Regardless, both capacity measures are greater than 0. This indicates super-capacity and thus facilitation among the two sub-processes. Curiously, this is at odds with the overadditivity of the RTs when both factors are in the low condition.

## DISCUSSION

Through the present results, we have demonstrated the ability of monkeys to perform a speeded response task with 2×2 factorial independent manipulations of difficulty. To our knowledge, this is a first application of these concepts in nonhuman primate research. We have also demonstrated the utility of systems factorial technology in assessing behavioral responses in such a way that can be used to infer underlying processing architectures. These pave the way for developing studies in monkeys that are directly comparable to studies in humans and extend investigation to the underlying neurophysiology producing the performance. We discuss two potential limitations of these current results: inter-monkey differences and error-prone performance. We conclude that neither of these considerations undermines the utility of this new experimental approach for nonhuman primate cognitive neurophysiology.

### Individual differences between monkeys

While we were able to identify the processing strategies used by the two monkeys, it was clear that the two monkeys used different strategies. This is notable for two reasons. First, it is important to note that subtle differences in RT behavior provide such different inferences for the mechanisms by which those RTs were generated. Second, these different inferences provide starkly contrasting predictions for the neurophysiological underpinnings of this behavior. In the case of Da, whose RTs suggest a parallel exhaustive processing architecture, neurons carrying signals related to the identifiability or the discriminability manipulations of the task may well be separate populations, or some neurons may only carry decision-related signals after both identifiability and discriminability decisions have been resolved. In the case of Le, whose RTs suggest a coactive processing architecture, there may not be separate populations of neurons sensitive to identifiability or discriminability, or there may indeed be a neural substrate that coalesces information relevant to both identifiability and discriminability.

Further, this dissociation between parallel exhaustive and coactive architectures has been described previously. Fifić and colleagues (2008a) had human participants perform a multidimensional classification task for stimuli whose dimensions were either separable or integral. Performance during classification of separable-dimension stimuli was marked by the use of a parallel exhaustive architecture whereas performance during classification of integral-dimension stimuli was marked by the use of a coactive architecture. This performance strategy difference, revealed only through systems factorial technology, is highly reminiscent of the performance strategy difference identified here. Because the shape and chromatic dimensions of the current stimuli are indeed different, they could be seen as separable dimensions. However, because both dimensions are carried by the same object they could be seen as integral. Monkey Da had performed several search tasks prior to this study, including tasks in which shape and color cue different aspects of the rules (e.g., Heitz et al., 2012; Reppert et al., 2018), which may help him separate these feature dimensions and thus use a parallel exhaustive strategy. Monkey Le, on the other hand, had not performed other tasks prior to the one in this study and thus may integrate the two feature dimensions and thus use a coactive strategy.

Many other investigators have addressed the problem of the architecture underlying visual search. All now agree that the slope of RT with set size is not an effective criterion. More complex tasks are needed. For example, previous work studying a wide variety of visual search displays with multiple targets concluded that whereas most search conditions are accomplished through parallel limited-capacity process, some limited conditions require serial search (Thornton & Gilden, 2007). A previous investigation of visual search with manipulation of target-distractor similarity employed systems factorial technology (Fifić et al. 2008b). These authors reported systematic departure from parallel or serial processing and concluded that the results were consistent with positively interacting parallel channels.

The particularities of doing sophisticated performance testing with macaque monkeys became evident also in the analysis of capacity, comparing RTs in blocks with 2×2 interleaved trial types with RTs in blocks with 2×1 or 1×2 interleaved trial types. The outcome of the analysis identified both monkeys working at super-capacity. While interest in how super-capacity can arise has been articulated (cf. Repperger et al., 2009), we should be mindful of trivial interpretations. Another interpretation of the appearance of super-capacity is that the monkeys applied different strategies to the 2×1 and 1×2 tasks because they were unusual in their training experience. This is evident by the longer RT in the 2×1 and 1×2 relative to the supposedly more difficulty 2×2 condition.

### Error-prone performance

Systems factorial technology generally assumes perfect or near-perfect performance, which is notable as errors can contaminate RT distributions through speed accuracy tradeoffs. However, performance was far from perfect in the data presented here. Thus, it is valid to wonder whether the SIC calculations and thus processing architecture inferences are mistaken due to contamination by errors. This seems unlikely for two reasons. First, the SIC curves for both monkeys are qualitatively similar to those obtained in several other studies involving systems factorial technology in humans with low error rates, and the resulting inferences are sensible in the context of separable and integral feature dimensions as discussed above. Second, simulation approaches that are allowed to produce errors have shown that the MIC and SIC signatures are robust with moderately high error rates (Fifić et al., 2008a; Townsend & Wenger, 2004). Specifically, coactive architectures tend to be the only architectures whose signatures degrade with errors by losing their indicative overadditivity. However, this means that a coactive architecture may be mistakenly labeled as serial exhaustive, so if an architecture is still identified as coactive in spite of high error rates this should only raise confidence in this inference. If anything, we suspect that the high error rate may have resulted in the uncharacteristic bimodality of the SIC curve for monkey Da, but the nature of this bimodality is not at odds with the overall inference of a parallel exhaustive architecture.

### Conclusions

RT in complex tasks is the summation of functionally distinct operations or stages. While not emphasized, the stage assumption is fundamental to the predominant model of “decision-making” – a single sequential-sampling process intervening between uninteresting visual encoding and response production stages. Such models explain performance and account for neural activity in visual discrimination tasks as well as visual search with direct stimulus-response mapping. But, if RT is not comprised of dissociable stages, then models like drift diffusion are disqualified and alternative models are endorsed, such as cascade (e.g., McClelland 1979) or asynchronous discrete flow (Miller 1988), which are qualitatively different mechanisms.

The most effective and perhaps only method for assessing the existence and characterizing the properties of modules or stages is the logic of separate modifiability. Crucially, single-stage decision-making models cannot explain tasks that require multiple, sequential operations. The term “decision” is hopelessly ambiguous when applied to a task that requires a “decision” about the location of a color singleton, a “decision” about the shape of the singleton, a “decision” about the shapes of distractors, a “decision” about the congruency of the singleton and distractor shapes, a “decision” about the instructed stimulus-response mapping, a “decision” about the correct endpoint of the saccade, and a “decision” about when to initiate the saccade. We have established that macaque monkeys can perform a task with simultaneous, independent factorial manipulations, producing performance measures that produce interpretable outcomes using the most advanced computational analytical approaches. This paves the way for a next step in cognitive neurophysiology of visual search.

## Acknowledgements

This work was supported by NIH RO1-EY08890, T32-EY007135, F32-EY028846, P30-EY008126, U54-HD083211 and by Robin and Richard Patton through the E. Bronson Ingram Chair in Neuroscience. We thank Greg Cox, Gordon Logan, Thomas Palmeri, and James Townsend for helpful comments and suggestions. Requests for materials should be addressed to JDS.

